# Multivariate empirical mode decomposition reveals markers of Alzheimer’s Disease in the oscillatory response to transcranial magnetic stimulation

**DOI:** 10.1101/2025.03.06.640967

**Authors:** Davide Bernardi, Elias P. Casula, Lorenzo Rocchi, Luciano Fadiga, Giacomo Koch, David Papo

## Abstract

**Objective:** To investigate EEG activity following transcranial magnetic stimulation (TMS) of the dorsolateral prefrontal cortex of Alzheimer’s Disease (AD) patients and control subjects using a data-driven characterization of brain oscillatory activity without prescribed frequency bands.

**Methods:** We employed multivariate empirical mode decomposition (MEMD) to analyze the TMS-EEG response of 38 AD patients and 21 control subjects. We used the distinct features of EEG oscillatory modes to train a classification algorithm, a support vector machine.

**Results:** AD patients exhibited a weakened slow-frequency response. Faster oscillatory modes displayed a biphasic response pattern in controls, characterized by an early increase followed by a widespread suppression, which was reduced in AD patients. Classification achieved robust discrimination performance (85%/23% true/false positive rate).

**Conclusions:** AD causes an impairment in the oscillatory response to TMS that has distinct features in different frequency ranges. These features uncovered by MEMD could serve as an effective EEG diagnostic marker.

**Significance:** Early detection of AD requires diagnostic tools that are both effective and accessible. Combining EEG with TMS shows great promise. Our results and method enhance TMS-EEG both as a practical diagnostic tool, and as a way to further our understanding of AD pathophysiology.

## 1. Introduction

The connection between Alzheimer’s Disease (AD), the most common cause of degenerative dementias (Gustavsson et al., 2023), and brain oscillations has been extensively investigated (Moretti et al., 2004; Yener and Başar, 2013; Babiloni et al., 2020, 2021; Ferreri et al., 2022). In particular, various studies have used EEG or MEG to characterize temporal or spectral differences in resting state brain activity between subjects with mild cognitive impairment (MCI) and AD dementia with respect to healthy controls (Jeong, 2004; Babiloni et al., 2009, 2010; Dauwels et al., 2010; Garcés et al., 2013; Wen et al., 2015; Babiloni et al., 2016; Blinowska et al., 2017; Vaghari et al., 2022). In general, MCI and AD patients are characterized by three important differences with respect to healthy control subjects (Cassani et al., 2018). First, slowing of resting oscillatory brain activity, thought to stem from loss of cholinergic innervation; in particular, resting electro- and magneto-encephalographic (EEG/MEG) activity in AD has mainly been associated with a decrease of fast oscillations over posterior brain regions and with a general enhancement of slow rhythms compared to healthy brain activity (Stam et al., 2002; Yener and Başar, 2013; Babiloni et al., 2020; Ferreri et al., 2022). Second, reduced synchrony, thought to reflect impaired neural communication pathways. Third, reduced signal complexity, possibly associated with neurodegeneration and reduced cortical connectivity. Furthermore, hyperexcitability of motor and sensory cortices has been documented (Ferreri et al., 2002; D’Amelio and Rossini, 2012), and a two-way relationship between changes in EEG/MEG rhythms and disease progression has been proposed (Stam et al., 2002; Iaccarino et al., 2016). Various studies also used EEG (Jelic et al., 2000; Rossini et al., 2006; Poil et al., 2013; Musaeus et al., 2020; Engedal et al., 2020; Rossini et al., 2022) and MEG (López et al., 2014) to predict progression from MCI to AD dementia.

Recently, EEG recordings have been paired with simultaneous transcranial magnetic stimulation (TMS), a noninvasive brain stimulation tool used to probe cortical excitability and functional connectivity (Ilmoniemi and Kičić, 2009; Rogasch and Fitzgerald, 2013; Siebner et al., 2022; Kallioniemi and Daskalakis, 2022; Hernandez-Pavon et al., 2023), to investigate physiological and pathological brain aging (Casarotto et al., 2011) not only in AD (Ferreri et al., 2016; Motta et al., 2018; Di Lazzaro et al., 2021; Casula et al., 2022a,b; Wu et al., 2022; Tăuţan et al., 2023), but also in other neurological diseases (Casula et al., 2024), (see also Nardone et al., 2020; Guerra et al., 2021; Ferreri et al., 2022; Farzan, 2024, for reviews).

The analysis of EEG oscillatory activity typically relies on predefined frequency bands, which offer a convenient approximation, particularly in clinical settings. However, the brain’s frequency spectrum is both content- and subject-specific, and traditional frequency band representations of intermittent signals can be challenging to interpret (Cohen, 1995; Huang et al., 1998; Mandic et al., 2013). To address these challenges, empirical mode decomposition (EMD) is a fully data-driven time-frequency analysis method that breaks down the original signal into simpler oscillatory components, called intrinsic mode functions (IMFs). Unlike traditional methods, which assume a fixed frequency structure, EMD is better suited for analyzing non-stationary signals—like those found in TMS-EEG recordings—where the signal characteristics change over time. Specifically, EEG signals modulated by TMS pulses often show frequency leakage, which can complicate interpretation with Fourier-based methods. EMD, in contrast, is well-suited for analyzing complex, non-periodic signals because it adapts to the signal’s variability and provides a clearer view of time-varying frequency components (Huang et al., 1998). While the initial EMD algorithm focused on univariate time series, multivariate extensions (Tanaka and Mandic, 2007; Rilling et al., 2007; Rehman and Mandic, 2010) facilitate the analysis of spatially extended brain dynamics captured by multielectrode EEG.

In this study, we employed multivariate empirical mode decomposition (MEMD) to investigate the amplitude modulation (AM) of EEG signals induced by single TMS pulses delivered to the left dorsolateral prefrontal cortex (DLPFC). This brain region has been identified as a clinical target in various pathologies, including depression (Sun et al., 2016; Voineskos et al., 2019), addiction (Gorelick et al., 2014), and pain (Moisset et al., 2016). Additionally, DLPFC appears to be one of the regions showing the most conspicuous changes in AD (Yener et al., 2007, 2008). Single-pulse TMS applied to the DLPFC elicits a local short-latency response (Wang et al., 2024; Gogulski et al., 2024b,a), which was suggested to reflect GABAergic inhibition and NMDA receptor function (Belardinelli et al., 2021), and associated with depression pathophysiology (Voineskos et al., 2019) and clinical response to TMS (Eshel et al., 2020).

Previous studies using prefrontal TMS stimulation focused on specific peaks in evoked potentials or the power within predefined frequency bands, finding some significant correlations with clinical parameters (Bagattini et al., 2019; Casula et al., 2022b; Asare et al., 2023). Notably, higher TMS-EEG evoked P30 amplitudes originating from the superior parietal cortex correlated with lower cognitive performance, as measured by MMSE and face–name memory scores. This augmentation was found to predict disease severity, suggesting an association between disease progression and effective long-distance fronto-parietal connectivity (Bagattini et al., 2019). Remarkably, later components (P50 and P70), elicited by TMS stimulation at dorsolateral prefrontal sites with different connectivity, exhibited no significant correlation with AD severity. This finding suggests that cognitive impairment at different stages of the disease is associated with specific neural circuits. Here, we explored whether MEMD could reveal novel features in the EEG response to TMS that are significantly altered in AD patients compared to a healthy control (HC) group. Importantly, the analysis we present here is applicable to any type of TMS-EEG dataset, and it has the potential to uncover distinct response patterns to TMS in other pathological brain conditions.

## 2. Methods

### 2.1. Experimental methods

#### 2.1.1. Subjects

The dataset was collected from 38 AD patients and 21 age-matched HC. AD patients screened for this study were recruited from the memory clinic of the Santa Lucia Foundation (Rome, Italy), where they presented with memory-related complaints. This study was approved by the review board and ethics committee of the Santa Lucia Foundation and was conducted following the principles of the Declaration of Helsinki and the International Conference on Harmonization Good Clinical Practice guidelines. All patients or their legal representatives provided written informed consent. Patients could withdraw at any point without prejudice. Patients were eligible if they had an established diagnosis of probable mild-to-moderate AD according to the International Working Group recommendations (Dubois et al., 2014). Inclusion criteria included: (1) age 50 to 85 years; (2) Clinical Dementia Rating Scale (CDR) score of 0.5-1; (3) Mini-Mental State Examination (MMSE) (Galasko et al., 1990) score of 18-26 at screening; (4) one caregiver; (5) treatment with acetylcholinesterase inhibitor for at least 6 months. Patients were excluded if they had extrapyramidal symptoms, history of stroke, other neurodegenerative disorders, psychotic disorders, or if they had been treated six months before enrollment with antipsychotic, antiparkinsonian, anticholinergic, and antiepileptic drugs. Signs of concomitant cerebrovascular disease on MRI scans were carefully investigated and excluded in all patients. Patients who agreed to participate underwent neuropsychological and neurophysiological evaluations. All technicians involved in the evaluations were blind to the patients’ characteristics. Age-matched HC were recruited after informed consent and underwent the same neurophysiological characterization as AD patients.

#### 2.1.2. Cognitive evaluation

Patients underwent the following cognitive and behavioral scales at enrollment and after 24 weeks: CDR Scale Sum of Boxes score (Morris, 1997); Alzheimer’s Disease Assessment Scale – Cognitive Subscale 11 (Fioravanti et al., 1994); Alzheimer’s Disease Cooperative Study - Activities of Daily Living (Galasko et al., 1997) and Neuropsychiatric Inventory (Cummings et al., 1994). These scales were chose based on their large use in studies assessing cognitive, behavioral and functional state in AD patients (Koch et al., 2022; Casula et al., 2022a,b).

#### 2.1.3. TMS-EEG protocol

During all recordings, participants sat on a comfortable armchair in a soundproof room in front of a computer screen. They were instructed to fixate a white cross (6 *×* 6 cm) in the screen center and keep their arms rested in a relaxed position. To avoid auditory event-related potentials, participants wore in-ear plugs that continuously played noise reproducing the time-varying frequencies of the TMS click (Massimini et al., 2005; Rocchi et al., 2021; Russo et al., 2022). The noise intensity was individually adjusted by increasing the volume (up to the maximum allowed by the equipment, which was 90 dB) until each participant confirmed that the TMS click was inaudible. TMS was performed using a Magstim Rapid2 magnetic biphasic stimulator connected to a figure-of-eight coil of 70 mm diameter. Coil positioning was guided by stereotaxic neuronavigation based on individual T1-weighted MRI volumes. To target the left DLPFC, the coil was positioned over the junction of the middle and anterior thirds of the middle frontal gyrus, corresponding to an area between the centre of the Brodmann area 9 and the border of Brodmann area 9 and Brodmann area 46. The coil orientation was set 45° away from the midline (Assogna et al., 2020; Casula et al., 2022b; Tăuţan et al., 2023). Stimulation intensity was based on a distance-adjusted motor threshold considering the individual coil-to-cortex distance (Stokes et al., 2007). The intensity of stimulation of single-pulse TMS was set at 90% of the distance-adjusted motor threshold. To ensure that this intensity was sufficient to evoke a reliable response in the AD patients, i.e. >40 V/m (Rosanova et al., 2009), we used a well-established procedure from our previous studies (Casula et al., 2022a,b; Koch et al., 2022; Tăuţan et al., 2023). First, we computed the scalp-to-cortex distance (SCD) and the induced E-field with SimNIBS v3.2, an open-source simulation package that integrates segmentation of MRI scans, mesh generation, and FEM E-field computations (Thielscher et al., 2015). For the HC group, we used MNI the standard brain (ERNIE) provided in SimNIBS software as an anatomical reference (Windhoff et al., 2013). To ensure that the two groups received the same stimulation in terms of intensity and efficacy, we compared the induced E-field. To induce a reliable cortical response, the single-pulse TMS intensity was 90% of the distance-adjusted motor threshold (Casula et al., 2022a; Maiella et al., 2022). TMS was delivered in 120 single-pulse blocks with random inter-stimulus interval in the 2-4 s range. EEG was recorded with a TMS-compatible DC amplifier (BrainAmp, BrainProducts GmbH, Munich, Germany) from 29 TMS-compatible Ag/AgCl pellet electrodes mounted on an elastic cap. Additional electrodes were used as a ground and reference. The ground was positioned in AFz, the reference on the nose tip. EEG signals were digitized with 5 kHz sampling rate. Skin/electrode impedance was maintained below 5 kΩ.

### 2.2. Data analysis

#### 2.2.1. Data preprocessing

TMS-EEG data were preprocessed offline with Brain Vision Analyzer (Brain Products GmbH, Munich, Germany). Data were segmented into 2s epochs centered on the TMS. TMS artifact was removed using a cubic interpolation, from 1 ms before to 10 ms following the pulse. We downsampled data to 1000 Hz and applied a 1-80 Hz Butterworth zero-phase band-pass and a 50-Hz notch filter. Visual inspection of all epochs revealed no apparent edge artifacts. Furthermore, the subsequent analysis focused on the central portion of each epoch. Following complete visual inspection, epochs with excessively noisy EEG were excluded from analysis (mean number of excluded epochs: 4.6*±*1.3). Independent component analysis was employed to identify and remove components reflecting muscle activity, eye movements, blink-related activity, and residual TMS artifacts based on previously established criteria (Maiella et al., 2022; Cristofari et al., 2023). The mean number of excluded components was 5.9*±*4.3. Finally, the signal was re-referenced to the average of all electrodes.

#### 2.2.2. IMF Extraction with noise-assisted MEMD

To extract IMFs, we used the noise-assisted MEMD (NA-MEMD) (Mandic et al., 2013), implemented adapting and optimizing a Python version^3^ of the original code^4^. We used 21 auxiliary noise channels with Gaussian random signals with prescribed power spectrum, generated as in ref. (Bernardi and Lindner, 2015; Bernardi et al., 2023). Their spectrum matched that of randomly selected channels. Noise amplitude was 10% of the signal. The stopping criterion for sifting was as in ref. (Rilling et al., 2003) with parameters: *θ*_1_ = 0.075 and *θ*_2_ = 10 *× θ*_1_ as threshold for global and local mean fluctuations, respectively; *α* = 0.075 as tolerance fraction.

We applied the NA-MEMD to each epoch, obtaining a variable number of IMFs per trial, from which we extracted instantaneous phase, amplitude, and frequency via Hilbert transform. To avoid unreliable IMFs and boundary artifacts (Mandic et al., 2013), we discarded IMFs outside the frequency range 5 Hz to 70 Hz and omitted the initial and final 250 ms. We tested the IMF reliability by analyzing sinusoidal mixtures with random phases and amplitudes plus auxiliary noise.

#### 2.2.3. IMF Classification

EEG data are inherently noisy, requiring averaging over trials for analysis. To classify IMFs across trials, we employed a two-step procedure: i) we defined three classification boundaries based on the minima of the histogram of all IMF average frequencies (fig. 2 dashed lines); ii) we applied a k-means clustering algorithm^5^ initialized using the boundaries from the previous step to adjust the classification to each subject.

**Figure 1:**
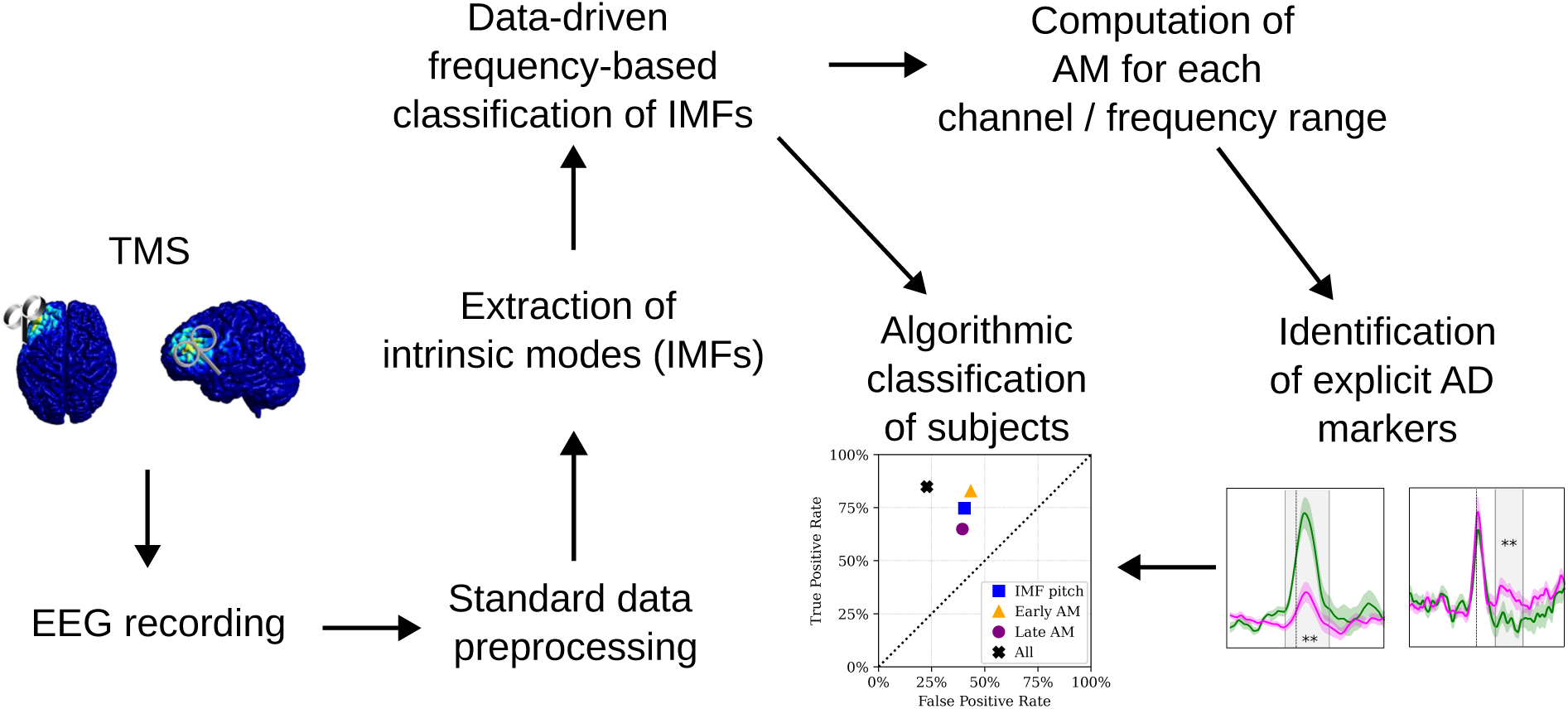
Graphical abstract of the analysis methods used in present study.

**Figure 2:**
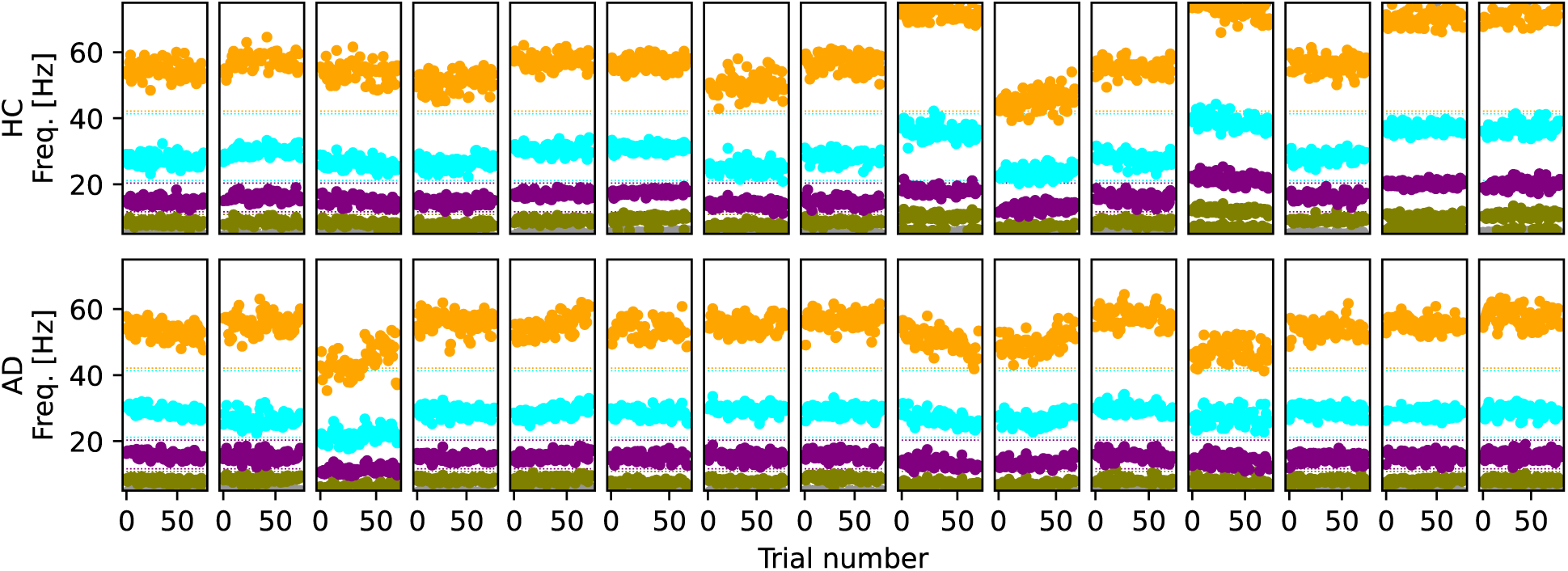
Subject-tailored IMF classification. Each panel corresponds to a subject, each dot represents the average frequency of one IMF per trial, color-coded based on the classification process detailed in the text. Colors indicate the frequency classification (olive: theta; purple: alpha; cyan: beta; orange: gamma). Each panel shows all IMF frequencies for one subject. In the top (bottom) row fifteen example subjects are chosen from the (HC) AD group.

Most trials revealed four IMFs, which clustered into four frequency ranges separated by gaps (see fig. 2), and they were consistently classified by the algorithm with a unique label. Starting from the lower frequency modes to the higher, we chose the labels IMF-1, IMF-2, IMF-3, and IMF-4.

Occasionally, fewer than four IMFs per trial were detected, indicating weak modes missed by the algorithm, which occurred in 0.9% of IMF-1, 0.6% of IMF-2, 0% of IMF-3, and 3.7% of IMF-4 trials. Mode splitting, where two IMFs shared the same label in a trial, was observed in slower frequencies (14.7% in IMF-1 range) and rarely in higher modes (0% in IMF-2, 0.1% in IMF-3, and 2.0% in IMF-4 ranges). In such cases, the IMF with slower frequency was chosen for further analysis.

### 2.2.4. Early and late AM response

The amplitude of each IMF was computed via Hilbert transform (Cohen, 1995). The trial-average of each IMF’s amplitude, ***w****_j,k,ℓ_*(*t*), was then computed for each frequency range *j* = 1, 2, 3, 4, channel *k*, and subject index *ℓ* = 1… 21 (HC) or *ℓ* = 1… 38 (AD). The standardized AM was defined as

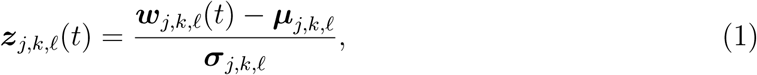

where ***µ****_j,k,ℓ_* and ***σ****_j,k,ℓ_* indicate the mean and standard deviation of ***w****_j,k,ℓ_*(*t*), respectively, computed using a time window terminating 100 ms before the TMS onset.

We compared the integrated AM response over two nonoverlapping time ranges. We defined the *early* response as

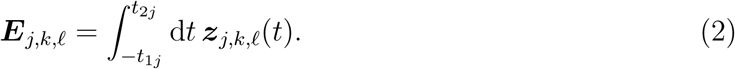

Note that, due to the Hilbert transform, the AM response is not causal and anticipates the signal, especially for slower frequencies. We defined as *late* response the integral

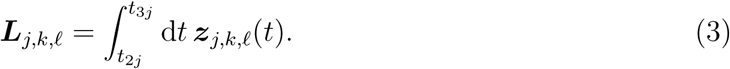

Parameters defining the time ranges were (in ms) *t*_11_ = 100*, t*_21_ = 350*, t*_31_ = 700; *t*_12_ = 100*, t*_22_ = 300*, t*_32_ = 650; *t*_13_ = 50*, t*_23_ = 120*, t*_33_ = 370; *t*_14_ = 50*, t*_24_ = 120*, t*_34_ = 370.

#### 2.2.5. Statistical analysis

We compared ***E****_j,k,ℓ_* for each *j, k* in the two conditions using a two-sided unequal variances Welch’s t-test. The same was done for ***L****_j,k,ℓ_*. Being ***E****_j,k,ℓ_* and ***L****_j,k,ℓ_* defined as integrals, they are expected to be normally distributed. The choice of this test, however, is not crucial, because the main purpose of the single-channel analysis was to identify spatial clusters at global level, which we did using the nonparametric cluster-based approach described by Maris and Oostenveld (2007). We defined spatial clusters as a set of channels fulfilling all following criteria: i) spatially contiguity; ii) difference in test statistic (t-value) with the same sign; iii) statistical significance at single-channel level (*p <* 0.05). Only clusters consisting of at least three channels were considered. To assess statistical significance at the cluster level, we constructed an empirical null distribution by evaluating the test statistic (the t-value of the entire cluster) across 10000 random partitions of the combined dataset. Based on this empirical null distribution, we computed a two-sided p-value for each cluster of channels. The cluster permutation test was implemented with custom Python code.

We performed a standard linear regression between the clinical scores mentioned in section 2.1.2 and the features used for the algorithmic classification described in the following subsection. Linear regressions and t-tests were performed using the open-source library scipy.stats.

#### 2.2.6. Subject Classification

We classified subjects by using a linear support vector machine, implemented using scikit-learn 1.1.2. The feature space was constructed using: the eight-dimensional space of the mean and dispersion of IMF frequencies; the projection of ***E****_j,k,l_* and ***L****_j,k,l_* onto the subspace of the three channels with the lowest p value smaller than 0.01. For cross-validation, the dataset was divided according to a 80%/20% ratio (training/test). Classification was repeated for 200 random data partitions. Since the two classes are unevenly represented, classification was repeated 100 times for each of the 200 data partitions by randomly oversampling the minority class during training and undersampling the majority class during testing.

## 3. Results

The baseline AD patients’ and HC characteristics are shown in table 1. No differences in sex, age, or education years were found between the two groups (all *p >* 0.05). The mean resting motor threshold (RMT) was 57.73*±*7.97 % maximum stimulator output (MSO) in the AD group and 64.11*±*9.49 % MSO in the HC group. There was a significantly lower RMT in the AD group (p=0.008). The mean SCD in the AD group was 18.7*±*1.7 mm. All participants received a stimulation >45 V/m (mean 57.7*±*3.8 V/m) with no differences between the mean E-field in the two groups (AD: 56.9*±*2.6; HC: 57.7*±*5.2; *p >* 0.05). The mean baseline MMSE score in AD patients was 17.84*±*5.86, which was significantly lower than in the HC group (28.4*±*2.06; *p <* 0.001).

**Table 1:** Comparison of AD and HC groups across various measures. *Indicates a significant p-value.

|  | Group | Mean | SD | p |
| --- | --- | --- | --- | --- |
| **Age (years)** | AD | 72.68 | 6.29 | 0.201 |
|  | HC | 70.2 | 8.06 |  |
| **Education (years)** | AD | 10.92 | 4.7 | 0.569 |
|  | HC | 10.2 | 4.68 |  |
| **Sex (% female)** | AD | 60.5 | - | 0.544 |
|  | HC | 52.3 | - |  |
| **MMSE** | AD | 17.84 | 5.86 | <0.001* |
|  | HC | 28.4 | 2.06 |  |
| **RMT** | AD | 57.73 | 7.97 | 0.008* |
|  | HC | 64.1 | 9.49 |  |
| **FAB** | AD | 10.83 | 3.79 | - |
|  | HC | - | - | - |
| **ADAS COG** | AD | 21.7 | 10.23 | - |
|  | HC | - | - | - |
| **NPI** | AD | 10.68 | 9.1 | - |
|  | HC | - | - | - |
| **ADCS - ADL** | AD | 57.39 | 12.48 | - |
|  | HC | - | - | - |
| **CDR** | AD | 4.53 | 2.51 | - |
|  | HC | - | - | - |

Before presenting our results for the IMF-related analysis, we briefly report on the bare TMS-evoked potentials to enable a visual assessment of the responses and overall signal quality. Figure 3 shows the TEPs recorded in four representative channels, as indicated in the respective inset. In channels near the stimulation site, a typical TEP response pattern with five peaks can be clearly observed, with a total TEP duration of approximately 250 ms.

**Figure 3:**
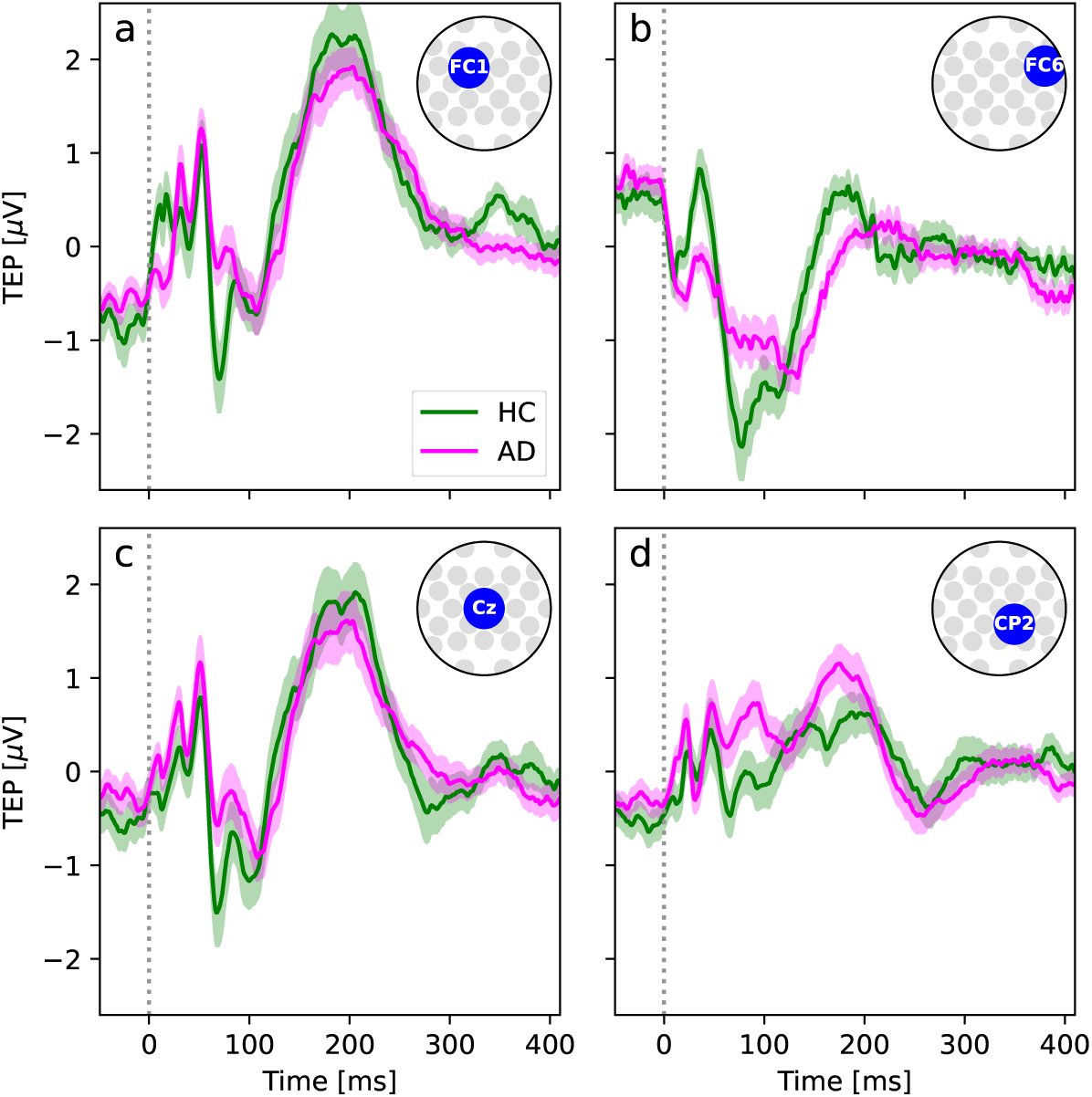
TMS-evoked potentials (TEPs) clearly display the usual five-peak oscillatory response. TEPs in the two conditions and the four selected channels shown in the corresponding inset. Shaded areas indicate the standard deviation of the mean.

### 3.1. Frequency alignment of IMFs with respect to classic bands

We now proceed to the analysis of IMFs obtained from MEMD. Before examining the TMS response, we will first illustrate how this method analyzes frequency content in a data-driven manner, adapting to each subject’s individual frequency profile. To this end, fig. 4 displays histograms of mean IMF frequencies found in four representative subjects, with each panel corresponding to one subject’s data. The frequency distributions of IMFs are grouped into four clusters labeled as IMF-1, IMF-2, IMF-3, and IMF-4 as described in the Methods. Each cluster is represented by a specific color (olive, purple, cyan, and orange, respectively). The classic EEG frequency bands within the range we considered for the IMF analysis (5-70 Hz) are theta (5-8 Hz), alpha (8-13 Hz), beta (13-30 Hz), and gamma (30-70 Hz). Boundaries between the classic bands are shown as vertical dotted lines, and the band labels are shown in black within the corresponding range. In each frequency histogram, segments that align with traditional bands are displayed in full color, while the portions outside these bands appear with increased transparency. For the two subjects shown in fig. 4a and b (AD subject 2 and HC subject 11, respectively), the IMF frequency distributions exhibit a strong alignment with the classic EEG bands, as the color-coded frequency clusters fall mostly within the established theta, alpha, beta, and gamma ranges. In contrast, fig. 4c and d show a marked deviation from this pattern, with IMF frequencies in these subjects (AD subject 3 and HC subject 10, respectively) not aligning within the traditional boundaries of the classic bands. Here, clusters extend outside the usual theta, alpha, beta, and gamma frequency ranges, suggesting that these subjects have frequency characteristics that do not conform to conventional EEG band patterns. This contrast exemplifies individual variability in frequency distributions, with some subjects adhering to typical band arrangements and others displaying unique frequency profiles, underscoring the value of data-driven approaches. Next, we proceed with the analysis of the AM in response to the TMS, following the pipeline sketched in fig. 1 and explained in the Methods, with the aim of finding novel potential markers of AD.

**Figure 4:**
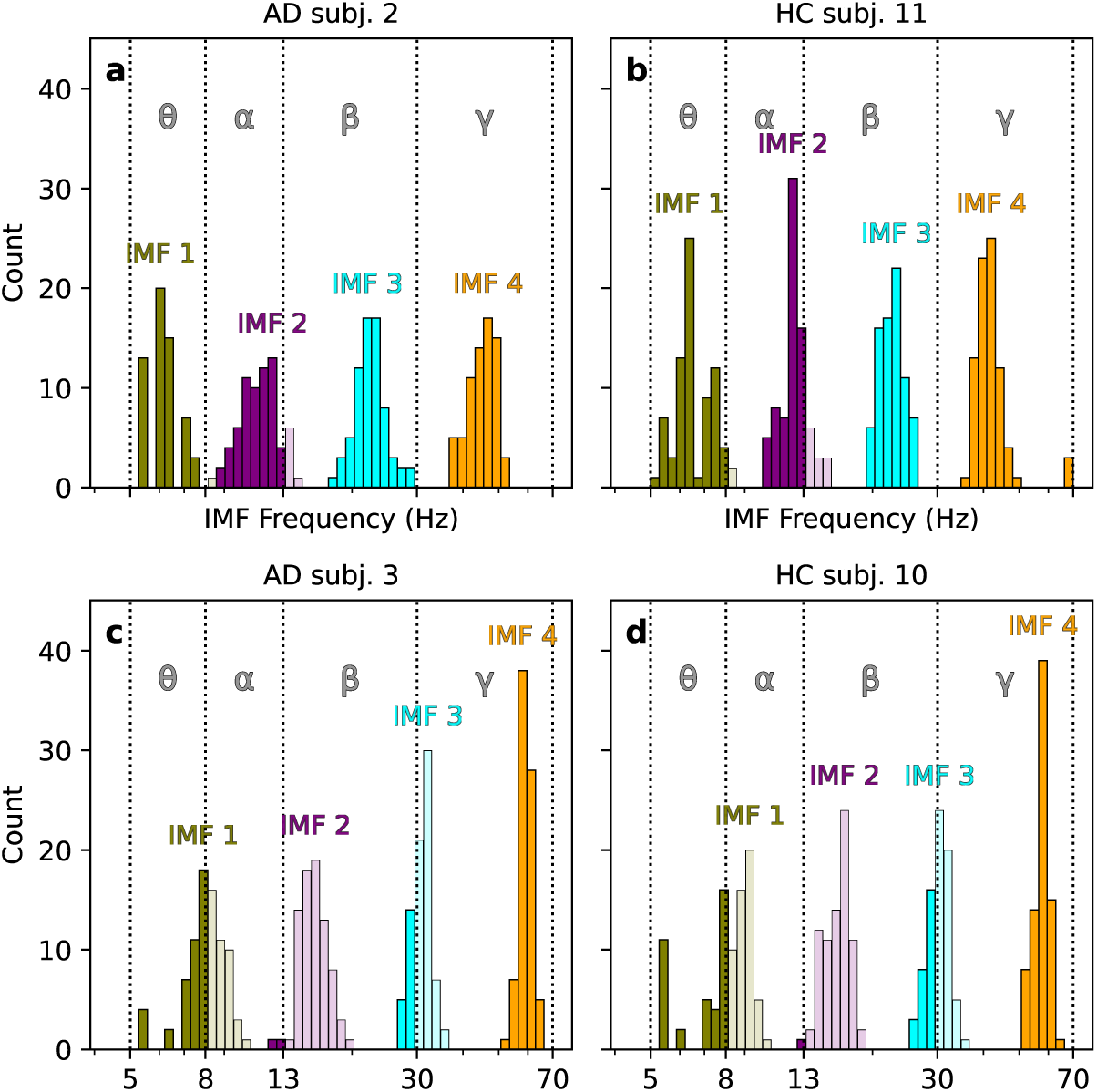
IMF frequencies do not necessarily align with standard frequency bands. Distribution of IMF frequencies for four representative subjects, classified as explained in the methods, and compared to the traditional frequency bands. For some subjects, such as AD subject 2 (a) and HC subject 11 (b), IMF frequencies primarily fall within the classic frequency bands, marked by vertical dotted lines and labeled with Greek letters. For others, like AD subject 3 (c) and HC subject 10 (d), IMF frequencies are less aligned with traditional bands. Segments of the frequency histogram that align with traditional bands are displayed in full color, while sections outside these bands appear with increased transparency.

### 3.2. Early AM modulation

We begin by discussing the early phase of the AM, i.e. the response of the oscillation amplitude of each IMF. For this purpose, we use Cz as an example channel. We chose this channel due to its neutral position. It is important to note that the purpose of the following single-channel discussion is solely to illustrate the shape of the AM response, as explained below. Therefore, the choice of this particular channel is not critical, as the response is consistent across most channels that show a strong signal.

We begin with the analysis of IMF-1, the slowest oscillatory mode we considered. We consistently observed a monophasic shape of the standardized oscillation amplitude *z*_1,Cz_(*t*) (see section 2.2.4c for the precise definition), i.e. it displayed a transient increase lasting about 400 ms (fig. 5a). In other words, the TMS elicited a temporary increase in the oscillation amplitudes of this mode in both conditions, which then returned to baseline.

**Figure 5:**
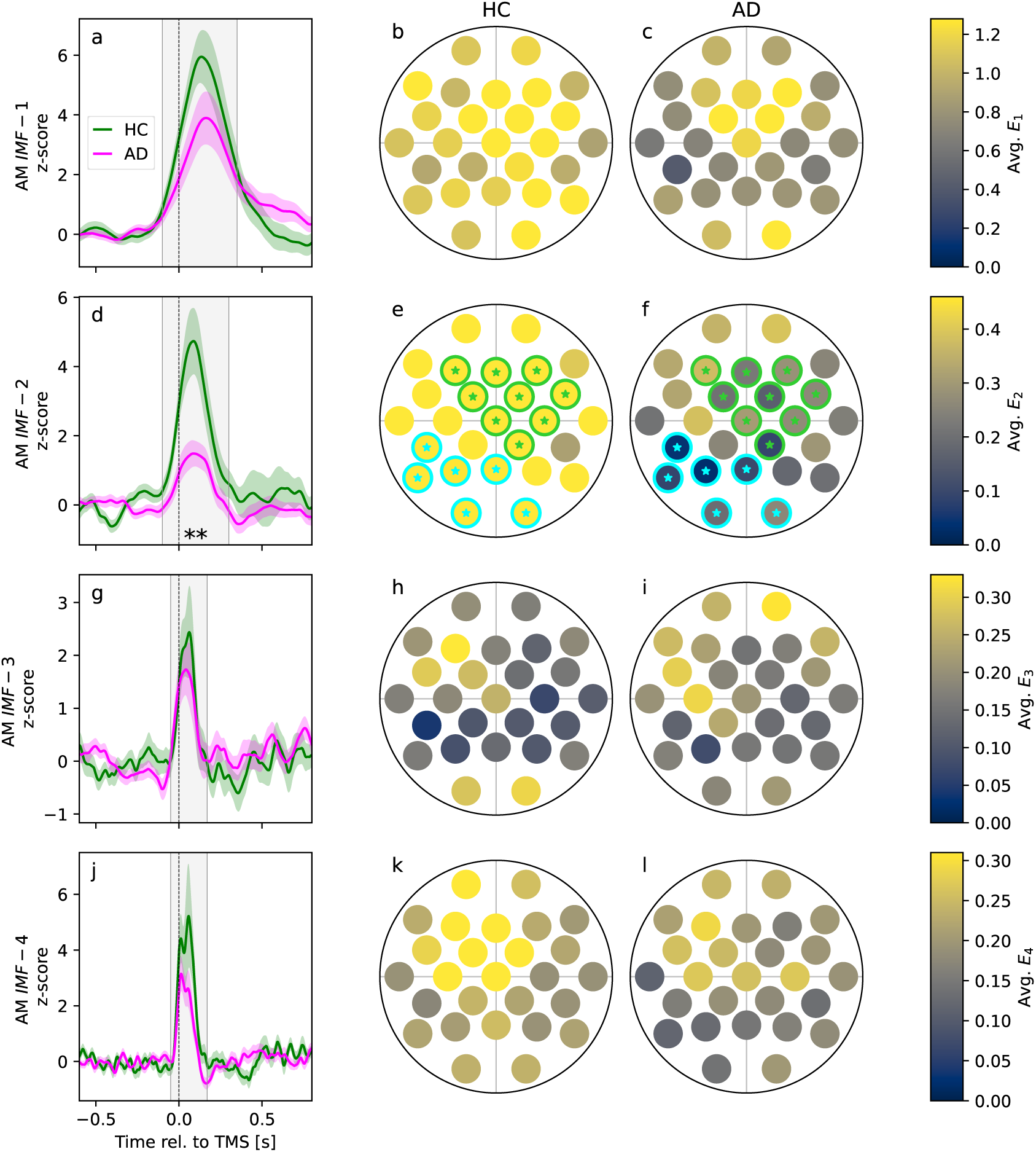
Early AM for slow IMF-2 is weaker in AD patients. (a) IMF-1 AM from channel Cz for HC (green) and AD patients (magenta); the gray shaded area is the time window used to compute the early response *E*_1_*_,k_*. (b,c): topographic representation of *E*_1_*_,k_* for HC and AD, respectively. (d-f), (g-i), (j-l): same for the IMF-2, IMF-3, and IMF-4, respectively. Highlighting and stars in the topographic representation of *E_j,k_* indicate significant channel clusters at the *p <* 0.05 level resulting from a nonparametric permutation test (see Methods).

To characterize the overall magnitude of the response to the pulse with a single number, we then considered the integrated early response ***E***_1,Cz_, averaged over subjects. The integrated early response corresponds to the area under the curve described by the oscillation amplitude within the shaded time window in fig. 5a (see the precise definition in section 2.2.4). There was no significant difference in this measure between the two conditions (***E***_1,Cz,HC_ = 1.79 *±* 0.29; ***E***_1,Cz,AD_ = 1.19 *±* 0.27; t-test, *p* = 0.14). Then, we compared the topographic distribution of ***E***_1_*_,k_* over all channels, shown in the topographic plot in fig. 5b,c. A clear response is seen in both conditions. The average response ***E***_1_*_,k_* appears stronger in HC in several channels, and, in two isolated channels, the difference was significant at the single-channel level (the full account of the ***E***_1_*_,k_* magnitude for each channel, IMF, and condition is given in table 2). However, we did not consider a significant difference at the single-channel level as sufficient evidence for a global difference in the two conditions. To perform this kind of assessment - taking both the need to correct for multiple comparisons and the spatial configuration into account - we employed the channel clustering procedure followed by the non-parametric permutation test (NPPT, see Methods section 2.2.5). This statistical analysis helps identify clusters of channels with significant differences by comparing actual data patterns to randomized data patterns. No significantly different cluster was detected.

**Table 2:** Results of t-test for the integrated early AM (see definition in section 2.2.4). Values in bold indicate significant p values at the single-channel level. A significant result at the single-channel level does not suffice for a significant difference between groups due to multiple comparisons. Neighboring significant channels were used to construct channel clusters for which a non-parametric permutation test was performed (see Methods section 2.2.5 for details).

| Channel | IMF 1 |  | IMF 2 |  | IMF 3 |  | IMF 4 |  |
| --- | --- | --- | --- | --- | --- | --- | --- | --- |
|  | t-score | p-value | t-score | p-value | t-score | p-value | t-score | p-value |
| Fp1 | 0.293 | 0.771 | 1.681 | 0.103 | -0.726 | 0.471 | 1.301 | 0.199 |
| Fp2 | 0.624 | 0.538 | 1.263 | 0.214 | -1.687 | 0.097 | 0.328 | 0.744 |
| F7 | 0.962 | 0.346 | 1.107 | 0.276 | -0.659 | 0.513 | 0.272 | 0.787 |
| F3 | -0.212 | 0.833 | 2.140 | **0.042** | 1.095 | 0.280 | 1.762 | 0.084 |
| Fz | 0.569 | 0.572 | 2.694 | **0.010** | 0.257 | 0.798 | 2.070 | **0.045** |
| F4 | 1.343 | 0.187 | 2.702 | **0.010** | -0.499 | 0.620 | 0.873 | 0.388 |
| F8 | 0.638 | 0.528 | 0.860 | 0.396 | -0.695 | 0.491 | 0.016 | 0.987 |
| FC5 | 1.437 | 0.157 | 1.511 | 0.139 | -0.058 | 0.954 | 0.313 | 0.756 |
| FC1 | 1.923 | 0.061 | 2.659 | **0.012** | 0.748 | 0.460 | 2.465 | **0.020** |
| FC2 | 1.361 | 0.179 | 3.696 | **0.001** | -0.177 | 0.861 | 1.739 | 0.092 |
| FC6 | 1.487 | 0.143 | 2.266 | **0.029** | -0.702 | 0.486 | 0.395 | 0.695 |
| T7 | 1.637 | 0.109 | 1.421 | 0.165 | -0.429 | 0.670 | 1.370 | 0.180 |
| C3 | 1.674 | 0.104 | 1.179 | 0.248 | -0.859 | 0.394 | 0.554 | 0.583 |
| Cz | 1.506 | 0.138 | 3.027 | **0.005** | 0.352 | 0.727 | 1.833 | 0.077 |
| C4 | 2.485 | **0.017** | 2.444 | **0.020** | -0.706 | 0.483 | -0.883 | 0.381 |
| T8 | 0.495 | 0.623 | 1.259 | 0.218 | -0.660 | 0.512 | -0.045 | 0.965 |
| CP5 | 1.873 | 0.069 | 2.300 | **0.027** | -1.049 | 0.299 | 0.251 | 0.803 |
| CP1 | 0.940 | 0.353 | 1.319 | 0.198 | -1.544 | 0.128 | 0.814 | 0.420 |
| CP2 | 1.143 | 0.260 | 2.776 | **0.010** | -0.359 | 0.721 | 0.570 | 0.571 |
| CP6 | 1.252 | 0.217 | 0.119 | 0.906 | -0.325 | 0.746 | 0.942 | 0.352 |
| P7 | 0.762 | 0.450 | 3.097 | **0.004** | -0.132 | 0.895 | 2.042 | **0.046** |
| P3 | 1.162 | 0.252 | 2.335 | **0.026** | 0.089 | 0.930 | 1.422 | 0.163 |
| Pz | 1.283 | 0.207 | 2.158 | **0.042** | -0.278 | 0.782 | 1.290 | 0.207 |
| P4 | 1.635 | 0.111 | 1.602 | 0.119 | -0.614 | 0.542 | 0.392 | 0.697 |
| P8 | 2.453 | **0.019** | 1.973 | 0.058 | 0.188 | 0.852 | 0.763 | 0.449 |
| O1 | 0.126 | 0.900 | 2.774 | **0.009** | 0.962 | 0.344 | 1.519 | 0.136 |
| O2 | -0.091 | 0.928 | 2.361 | **0.025** | 0.934 | 0.355 | 0.580 | 0.564 |

In the IMF-2 range, we observed a similar monophasic shape in the AM (fig. 5d). Here, however, the difference between the conditions was more pronounced (***E***_2,Cz,HC_ = 1.01 *±* 0.21; ***E***_2,Cz,AD_ = 0.31 *±* 0.10, *p* = 0.005, t-test), indicating a significantly weaker response in AD patients. Applying the channel clustering and the NPPT revealed two channel clusters that differed significantly, with a stronger response in HC. One cluster was located in the right frontal area (fig. 5e,f, green markings, *p* = 0.001, NPPT), the other cluster was located in the left parietal area (cyan markings, *p* = 0.0003, NPPT).

For the faster IMF-3, the AM was similar in the two conditions (fig. 5g; ***E***_3,Cz,HC_ = 0.25 *±* 0.10; ***E***_3,Cz,AD_ = 0.21 *±* 0.08). The topographic plot (fig. 5h,i) shows that the response was localized mostly in central areas. No significantly different channel cluster was detected. In the IMF-4 range (the fastest oscillation mode we considered), we observed a secondary peak in *z*_4,Cz_(*t*) about 50 ms after the first peak (fig. 5j) (***E***_4,Cz,HC_ = 0.52 *±* 0.13; ***E***_4,Cz,AD_ = 0.26 *±* 0.06). This peak contributed to a mild increase of ***E***_4_*_,k_* in several central electrodes in HC (fig. 5k,l). However, no statistically significant channel cluster was found.

Overall, these results suggest that while the early AM response shows transient increases in both conditions, the magnitude and pattern vary, especially in IMF-2, where the HC group displays a significantly stronger response than the AD group.

### 3.3. Late AM modulation

As reported below, the late response displayed the most notable effects in the contralateral frontal and parietal areas. Therefore, to illustrate the shape of the late AM response, we use FC2 as an example channel. As for the previous analysis, the choice of the specific channel is only for illustration purposes and is not relevant for the results.

For the slowest IMF-1, the late AM response was similar between the two conditions (fig. 6a). Essentially, there was no late response in both groups, as, within the relevant time window (gray shaded area in fig. 6a) the oscillation amplitude was at baseline level in both experimental groups. Hence, also the measure quantifying the integrated late response ***L***_1,FC2_, defined in eq. (3) as the area below the curve, was small for both conditions (***L***_1,FC2,HC_ = 0.14 *±* 0.11; ***L***_1,FC2,AD_ = 0.26 *±* 0.12). At the global level, no significant difference between the two conditions was observed (fig. 6b, c). The full account of the ***L*** magnitude for each channel, IMF, and condition is given in table 3.

**Figure 6:**
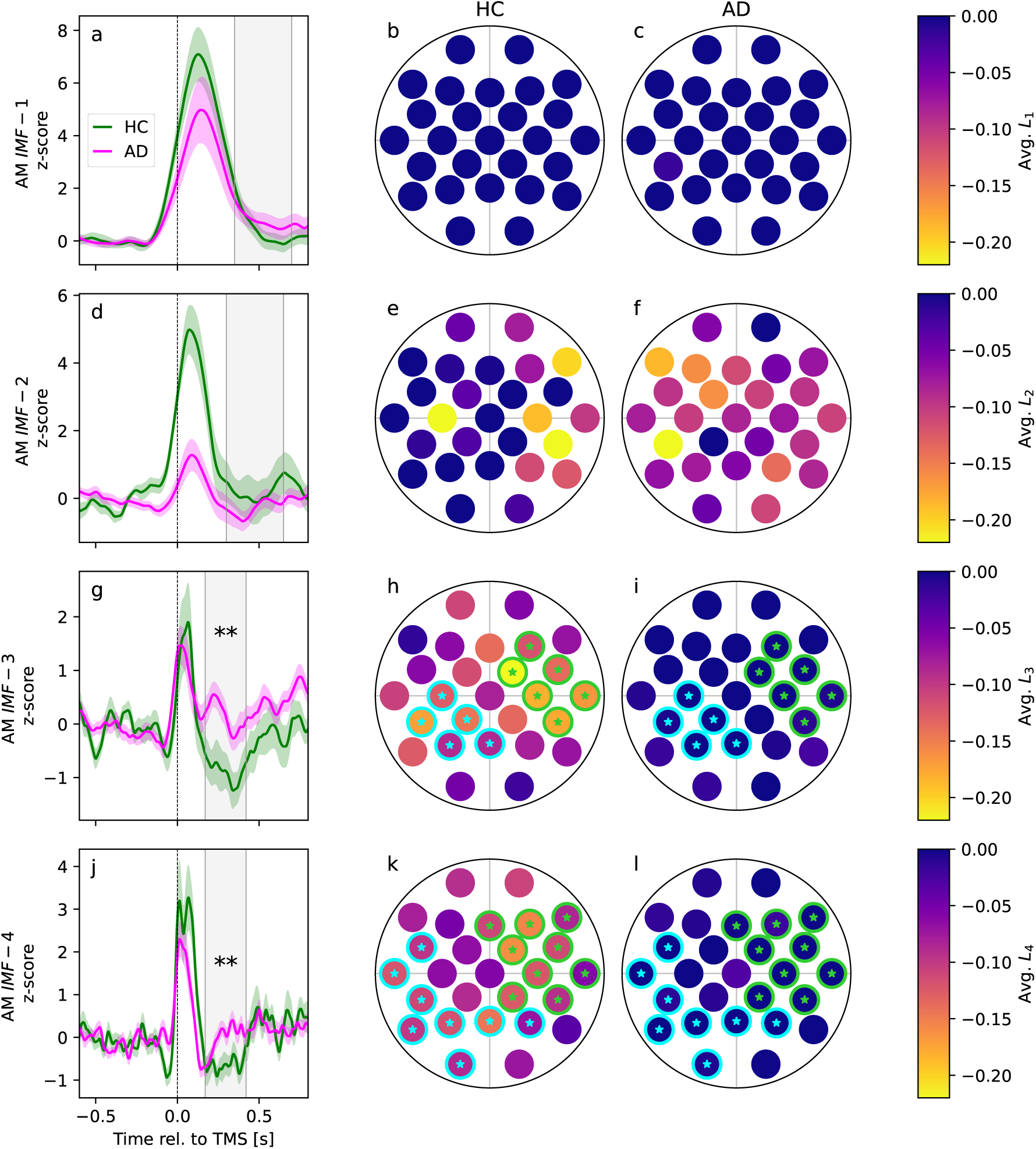
Late IMF-3 and IMF-4 response show a transient oscillation suppression in HC, which is absent in AD patients. (a) IMF-1 AM from channel FC2 for HC (green) and AD patients (magenta); the gray shaded area is the time window used to compute the late response *L*_2_*_,k_*. (b,c): topographic representation of *E*_2_*_,k_* for HC and AD, respectively. (d-f), (g-i), (j-l): same for IMF-2, IMF-3, and IMF-4, respectively. Highlighting and stars indicate significant channel clusters at the *p <* 0.05 level resulting from a nonparametric permutation test (see Methods).

**Table 3:** Results of t-test for the integrated late AM (see definition in section 2.2.4).Values in bold indicate significant p values at the single-channel level. A significant result at the single-channel level does not suffice for a significant difference between groups due to multiple comparisons. Neighboring significant channels were used to construct channel clusters for which a non-parametric permutation test was performed (see Methods section 2.2.5 for details).

| Channel | IMF 1 |  | IMF 2 |  | IMF 3 |  | IMF 4 |  |
| --- | --- | --- | --- | --- | --- | --- | --- | --- |
|  | t-score | p-value | t-score | p-value | t-score | p-value | t-score | p-value |
| Fp1 | -0.543 | 0.589 | 0.130 | 0.897 | -1.324 | 0.193 | -2.075 | **0.044** |
| Fp2 | -0.873 | 0.388 | -0.594 | 0.556 | -1.093 | 0.280 | -2.703 | **0.009** |
| F7 | -0.442 | 0.661 | 2.000 | 0.053 | -0.302 | 0.764 | -1.403 | 0.169 |
| F3 | -0.984 | 0.330 | 1.069 | 0.293 | -0.797 | 0.430 | -1.140 | 0.260 |
| Fz | -0.946 | 0.349 | 0.734 | 0.469 | -1.939 | 0.059 | -2.087 | **0.044** |
| F4 | -0.443 | 0.661 | -0.135 | 0.893 | -2.098 | **0.042** | -3.106 | **0.003** |
| F8 | -1.804 | 0.077 | -0.818 | 0.420 | -1.547 | 0.129 | -2.753 | **0.008** |
| FC5 | 0.036 | 0.971 | 1.219 | 0.234 | -1.646 | 0.106 | -2.709 | **0.009** |
| FC1 | -0.591 | 0.559 | 0.816 | 0.421 | -1.441 | 0.159 | -1.602 | 0.118 |
| FC2 | -0.721 | 0.474 | 0.849 | 0.403 | -2.972 | **0.005** | -3.042 | **0.004** |
| FC6 | -0.083 | 0.935 | 0.443 | 0.662 | -3.239 | **0.002** | -3.460 | **0.001** |
| T7 | -0.065 | 0.949 | 0.637 | 0.530 | -1.595 | 0.119 | -2.666 | **0.011** |
| C3 | -0.719 | 0.478 | -1.141 | 0.262 | -2.548 | **0.015** | -1.799 | 0.078 |
| Cz | -1.080 | 0.286 | 0.890 | 0.381 | -1.274 | 0.210 | -0.464 | 0.645 |
| C4 | -0.879 | 0.384 | -0.865 | 0.393 | -2.797 | **0.008** | -2.842 | **0.006** |
| T8 | -1.087 | 0.285 | 0.046 | 0.964 | -2.404 | **0.020** | -2.025 | **0.048** |
| CP5 | 0.348 | 0.730 | 0.935 | 0.359 | -2.263 | **0.030** | -2.645 | **0.011** |
| CP1 | -0.050 | 0.960 | -0.135 | 0.894 | -3.211 | **0.003** | -1.424 | 0.161 |
| CP2 | 0.230 | 0.820 | 0.284 | 0.779 | -1.854 | 0.072 | -2.469 | **0.018** |
| CP6 | 0.608 | 0.548 | -1.064 | 0.293 | -2.201 | **0.034** | -2.450 | **0.018** |
| P7 | 0.302 | 0.764 | 0.755 | 0.458 | -1.633 | 0.110 | -2.219 | **0.032** |
| P3 | 0.012 | 0.990 | 0.706 | 0.486 | -2.288 | **0.028** | -2.925 | **0.005** |
| Pz | 0.334 | 0.741 | 0.850 | 0.405 | -2.107 | **0.043** | -3.101 | **0.003** |
| P4 | 0.938 | 0.357 | 0.197 | 0.844 | -1.006 | 0.323 | -2.358 | **0.022** |
| P8 | 0.149 | 0.883 | -0.299 | 0.766 | -0.769 | 0.446 | -1.560 | 0.124 |
| O1 | -0.905 | 0.373 | 0.653 | 0.521 | -0.465 | 0.645 | -2.126 | **0.041** |
| O2 | -0.411 | 0.684 | 0.565 | 0.577 | -0.555 | 0.582 | -1.781 | 0.082 |

In the case of IMF-2, we also did not observe a significant difference in the late AM response between conditions (fig. 6d ***L***_2,FC2,HC_ = 0.06 *±* 0.18; ***L***_2,FC2,AD_ = *−*0.11 *±* 0.07). The global analysis of ***L***_2_*_,k_* (fig. 6e,f) showed no consistent pattern or significantly different clusters, further suggesting minimal variation across channels in this oscillatory mode.

For faster oscillatory modes, significant differences emerged between the two conditions. Specifically, in HC, the amplitude of IMF-3 showed a notable suppression lasting several hundred milliseconds, beginning approximately 100 ms after the TMS pulse (fig. 6g). The AM response in HC took on a biphasic shape, with an initial peak followed by a late “rebound" suppression. In contrast, this late suppression was nearly absent in AD patients, leading to a statistically significant difference between the groups at the single-channel level (***L***_3,FC2,HC_ = *−*0.23 *±* 0.07; ***L***_3,FC2,AD_ = 0.04 *±* 0.05; *p* = 0.002, t-test). To assess whether there was a significant difference between groups at the global level, we applied the clustering procedure followed by the NPPT. We found two channel clusters, in which ***L***_3_*_,k_* was significantly more negative in HC than in AD patients, where it was nearly zero. One cluster was located within the controlateral frontal area (fig. 6h,i, green markings *p* = 0.002, NPPT), while the other was located in the left and central parietal areas (fig. 6h,i, cyan markings *p* = 0.002, NPPT).

For IMF-4, we observed a similar general picture. A late negative response was present in HC, but not in AD patients (fig. 6j, ***L***_4,FC2,HC_ = *−*0.16 *±* 0.04; ***L***_4,FC2,AD_ = 0.00 *±* 0.03, *p* = 0.004, t-test). This late suppression, seen in most channels of the right frontal and parietal areas in HC (fig. 6k), was nearly absent in AD patients (fig. 6l). We found two large clusters located within the frontal right lobe (fig. 6k-l, green cluster) and in the left and central parietal areas (fig. 6k-l, cyan cluster), respectively, in which this difference was significant (*p* = 0.0005 and *p* = 0.0003, respectively, NPPT).

Overall, these findings indicate that for slower oscillatory modes (IMF-1 and IMF-2), there were minimal late AM responses and no significant differences between the two conditions. However, faster oscillatory modes (IMF-3 and IMF-4) revealed significant group differences: in the HC group, there was a marked suppression that was largely absent in AD patients. This difference was observed in clusters located in contralateral frontal and parietal areas.

### 3.4. Subject algorithmic classification

Next, we explored whether IMF features could be used to classify subjects individually. As detailed in the Methods section, we used a standard classification algorithm trained on the IMF features discussed above. We evaluated the classification performance using a standard Receiver Operating Characteristic (ROC) analysis (fig. 7), which is a common method to assess the accuracy of a classifier. In an ROC analysis, the true positive rate (TPR) is plotted versus the false positive rate (FPR). The TPR, also called sensitivity, indicates how often the classifier correctly identifies positive cases. The FPR, which is linked to specificity, shows how often the classifier incorrectly identifies a negative case as positive. Specificity itself is calculated as 1 *−* FPR. In the ROC plot, the top left corner represents ideal performance, where the classifier has a perfect true positive rate (100%) and a zero false positive rate. The diagonal line represents a performance equal to random chance, where the classifier has an equal probability of identifying true and false positives.

**Figure 7:**
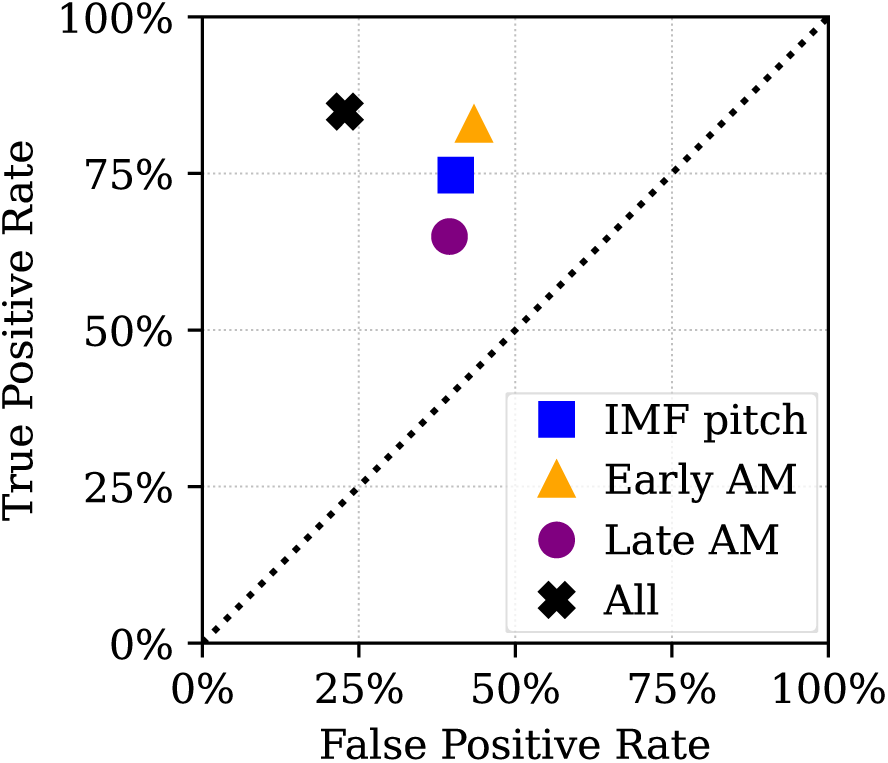
Successful subject classification based on the TMS-EEG markers found via the MEMD analysis pipeline (see fig. 1). Receiver Operating Characteristic analysis for the subject classification using a support vector machine trained on the IMF features as indicated in the legend and described in the main text.

We first trained the classifier using each feature individually, to probe the power of each single feature. Then, we trained the classifier using all features together, to check if features were redundant or if they contributed in part uniquely to classification performance (see Methods for details).

When we trained the classifier using only the IMF frequencies, which reflect spontaneous activity, the classifier achieved a false positive rate (FPR) of 40% and a true positive rate (TPR) of 75% (fig. 7, blue square). Training the classifier using only the early AM response ***E*** increased FPR and TPR to 43% and 83%, respectively (orange triangle), i.e. it had a higher sensitivity but a lower specificity. Training the classifier on the late response ***L*** decreased FPR and TPR to 39% and 65%, respectively (purple circle), i.e. it decreased the sensitivity but it increases the specificity. Finally, combining all features led to a consistent improvement in classification performance, reducing the FPR to 23% and increasing the TPR to 85% (black cross), thus improving both sensitivity and specificity. This improvement observed when combining all features suggests that features were not redundant, contributing independently to the classifier’s performance.

### 3.5. No significant linear correlation of early or late AM with clinical scores

To assess whether a simple linear relationship existed between the features used for algorithmic classification and disease severity, we conducted a linear regression analysis between each clinical score and the subject-related metrics extracted from the TMS-EEG responses, and used for the algorithmic classification described in the previous subsection.

A detailed report is provided in table 4. After applying the Holm-Bonferroni correction for multiple comparisons, no significant correlations were found.

**Table 4:** Linear regression results for Early and Late AM across Baseline and 24 week follow-up. Note that due to the correction for multiple comparisons, there is no statistically significant correlation.

| Clinical Score | Time | Early AM |  | Late AM |  |
| --- | --- | --- | --- | --- | --- |
| | | $r$ | $p$ | $r$ | $p$ |
| ADAS PC | Baseline | 0.251 | 0.153 | -0.338 | 0.050 |
|  | 24 Weeks | 0.056 | 0.800 | -0.157 | 0.475 |
| ADAS PG | Baseline | 0.231 | 0.189 | -0.357 | 0.038 |
|  | 24 Weeks | 0.046 | 0.836 | -0.195 | 0.373 |
| ADCS ADL | Baseline | -0.119 | 0.483 | -0.149 | 0.377 |
|  | 24 Weeks | -0.065 | 0.738 | -0.152 | 0.431 |
| CDR | Baseline | 0.105 | 0.536 | -0.070 | 0.683 |
|  | 24 Weeks | 0.004 | 0.985 | 0.122 | 0.529 |
| FAB PC | Baseline | -0.279 | 0.105 | 0.328 | 0.054 |
|  | 24 Weeks | -0.231 | 0.277 | 0.160 | 0.456 |
| FAB PG | Baseline | -0.246 | 0.154 | 0.358 | 0.034 |
|  | 24 Weeks | -0.201 | 0.347 | 0.238 | 0.262 |
| MMSE PC | Baseline | -0.172 | 0.309 | 0.250 | 0.136 |
|  | 24 Weeks | -0.209 | 0.328 | 0.158 | 0.462 |
| MMSE PG | Baseline | -0.185 | 0.273 | 0.265 | 0.113 |
|  | 24 Weeks | -0.168 | 0.432 | 0.185 | 0.388 |
| NPI | Baseline | 0.296 | 0.076 | 0.153 | 0.368 |
|  | 24 Weeks | 0.103 | 0.594 | 0.064 | 0.740 |

## 4. Discussion

Using MEMD, we analyzed the EEG response to TMS targeting the left DLPFC, identifying IMFs spanning various frequency ranges with considerable individual variability. Our results revealed an initial increase in oscillatory amplitude post-TMS across most frequency ranges in both HC and AD patients. However, two key distinctions emerged between the groups. Firstly, HC exhibited significantly stronger early response amplitudes in IMF-2. Secondly, a biphasic response characterized by a transient suppression of oscillation amplitude in IMF-3 and IMF-4 was prominent in HC, but notably reduced in AD patients. This discrepancy was localized to two electrode clusters, one in the contralateral prefrontal lobe and the other in parietal areas.

No significant correlations were observed between the magnitude of early and late responses. This, along with the spatial and temporal separation, suggests distinct underlying neurobiological mechanisms. Given the complexity and parameter-specific nature of the brain’s response to TMS (Siebner et al., 2022), it is plausible that multiple neurobiological mechanisms contribute to the early and late responses rather than a singular cause. A prevalent hypothesis proposes that AD disrupts the excitatory-to-inhibitory (E/I) balance in vulnerable brain regions and neuron types (Palop et al., 2007; Verret et al., 2012; Limon et al., 2012; Busche and Konnerth, 2016; Styr and Slutsky, 2018). This imbalance is attributed to GABAergic neuron dysfunction, potentially serving as a mechanism underlying hyperactivity and hypersynchrony of pyramidal neurons (Ramírez-Toraño et al., 2020; Lauterborn et al., 2021). Neuronal circuits require recurrent inhibition to respond to stimuli in a time-locked manner (Haider et al., 2013; Lefebvre et al., 2015; Kühn and Helias, 2017). Thus, weakened inhibition can result in an impairment of the oscillatory response, as observed in our analysis of the early response. Regarding the late response, the transient suppression in higher oscillatory frequencies implies a temporary desynchronization of these rhythms. Generally, it has been observed in cortical networks that external input can dampen spontaneous oscillations (Tan et al., 2014; Stringer et al., 2016), and experimental as well computational studies highlighted the role of feedforward and recurrent GABAergic inhibition in suppressing oscillations and signal gating (Vogels and Abbott, 2009; Renart et al., 2010; Sippy and Yuste, 2013; Helias et al., 2014; Bernardi and Lindner, 2019; Bernardi et al., 2021). Furthermore, the time delay of the late response we observed aligns with the kinetics of slow NMDA receptors and multisynaptic relays. Importantly, recent evidence from paired TMS and intracranial EEG in epilepsy patients shows that TMS applied to the DLPFC elicits not only a local response but also a response in distant, functionally connected brain regions after a time delay similar to the late response suppression observed here (Wang et al., 2024). Therefore, it is plausible that the reduced functional connectivity in AD patients (López-Sanz et al., 2017; Pusil et al., 2019; Chino et al., 2022) contributes to the observed absence of the late response. Both our main observations can thus be reconciled with the E/I imbalance hypothesis and, more broadly, with a system that is less responsive to external stimuli.

The significance of our results was further corroborated by the robust classification performance in discriminating AD patients from HC achieved by a classification algorithm trained using spontaneous and evoked IMF features. Importantly, our approach avoids the ‘black-box’ issue of brute-force machine learning algorithms, because we trained the algorithm on explicit and neurobiologically interpretable features.

Finally, neither early nor late AM patterns significantly correlated with scores in a variety of clinical tests. This lack of correlation does not directly contradict previous mixed reports of a significant negative correlation between P30 and cognitive decline severity as measured by the MMSE (Bagattini et al., 2019) or of a positive correlation with the Neuropsychiatric Inventory Questionnaire severity score (Asare et al., 2023), because the amplitude of specific peaks in the evoked potential cannot be directly related to the AM of any specific IMF. The absence of direct correlations with clinical scores suggests that the effects observed here might offer a distinct marker of AD, independent of traditional cognitive tests. This independence could potentially enhance their diagnostic usefulness.

The use of MEMD, or even basic EMD, in the TMS-EEG context has generally been limited, and to our knowledge, there has been no prior application to AD patients or other neuropathologies. One previous study applied EMD to a TMS-EEG dataset to detect the onset of TMS-evoked potentials in synthetic data and tested it on a small group of subjects (Pigorini et al., 2011), though they subsequently employed a different method based on the Hilbert-Huang Spectrum. Another preliminary investigation explored combining MEMD with source localization in simulations and data from a single subject (Hansen et al., 2019). Thus, the current approach is novel and can be applied to any TMS-EEG dataset, potentially revealing unique response patterns to TMS in various pathological brain conditions.

## 5. Conclusions

Our study revealed features of the oscillatory response to TMS that could serve as markers for distinguishing AD patients from HC, with the potential of offering a fast and minimally invasive diagnostic approach. From a methodological viewpoint, our findings highlight the versatility of adaptive data-driven algorithms, demonstrating their compatibility with trial-averaging while maintaining connections with traditional methods in EEG time-frequency analysis.

## Acknowledgments

This work was supported by the Italian Ministry of University and Research under the National Recovery and Resilience Plan (Fondi DM 502/2022 – PNRR MC42 Bando Giovani Ricercatori) to EPC. This work has been partially funded by PRIN 2020 to LF.

## Author contributions

DB: Conceptualization, Methodology, Software, Validation, Formal analysis, Data Curation, Visualization, Writing - Original draft, Writing - Review & editing. EPC: Conceptualization, Software, Investigation, Data Curation, Writing - Original draft, Writing - Review & editing. LR: Data Curation, Resources. LF: Conceptualization, Resources, Supervision, Writing - Review & editing, Project administration, Funding acquisition. GK: Conceptualization, Resources, Supervision, Writing - Review & editing, Project administration, Funding acquisition. DP: Conceptualization, Methodology, Supervision, Writing - Original draft, Writing - Review & editing, Project administration.

## Abbreviations

AD: Alzheimer’s Disease
ADAS - COG: Alzheimer’s Disease Assessment Scale - Cognitive Subscale
ADCS - ADL: Alzheimer’s Disease Cooperative Study - Activities of Daily Living
AM: Amplitude Modulation
CDR: Clinical Dementia Rating
DLPFC: Dorsolateral Prefrontal Cortex
EEG: Electroencephalography
FAB: Frontal Assessment Battery
HC: Healthy Control
IMF: Intrinsic Mode Function
MEG: Magnetoencephalography
MMSE: Mini-Mental State Examination
MEMD: Multivariate Empirical Mode Decomposition
MSO: Maximum Stimulator Output
MMSE: Mini-Mental State Examination
NPPT: Non-parametric permutation test
NPI: Neuropsychiatric Inventory
RMT: Resting Motor Threshold
SCD: Scalp-to-cortex Distance
TEP: TMS-evoked potential
TMS: Transcranial Magnetic Stimulation

## Footnotes

3 https://github.com/mariogrune/MEMD-Python

4 https://www.commsp.ee.ic.ac.uk/~mandic/research/emd.htm

5 scikit-learn 1.1.2 (Pedregosa et al., 2011)

